# Analytical expectations for ancestry junction accumulation in admixed genomes

**DOI:** 10.1101/2025.10.28.685223

**Authors:** Shirin Nataneli, Aydin Loid Karatas, Tessa Ferrari, Roshni A. Patel, Jazlyn A. Mooney

**Affiliations:** Department of Quantitative and Computational Biology, University of Southern California, Los Angeles, CA, USA; Department of Data Science, University of Oregon, Eugene, Oregon, USA; Institute of Ecology and Evolution, University of Oregon, Eugene, Oregon, USA

## Abstract

Complex demographic events have shaped human history and genetic variation across the genome. Here, we investigate the recent evolutionary history of admixed populations that descend from distinct ancestral sources. We present a discrete, generalizable model of admixture that leverages ancestry switches, which are recombination breakpoints that mark changes in ancestral origin along a chromosome. We derive analytical expectations for the number of ancestry switches within a genomic segment as functions of recombination rate, ancestry heterozygosity, and effective population size. We then extend these expectations to incorporate population-specific recombination maps. Our theoretical predictions are in close agreement with forward-in-time simulations that we use to trace ancestry junction accumulation since an initial admixture event with both constant and variable recombination models. We observe minimal variability in switch counts across ten simulation replicates, underscoring the robustness of the theoretical expectation. Furthermore, model-based switch counts, parameterized using literature-informed demographic values, agree with empirical observations from African American individuals in the 1000 Genomes Project. For example, when modeling human chromosome 1, we found a mean of approximately six switches per haplotype, which aligns with the theoretical expectation under an initial African ancestry proportion of 0.85, and agrees with published estimates from other African-American cohorts. Overall, the model provides a new route for using ancestry switches to understand how recombination and demography jointly shape ancestry patterns in admixed populations without requiring separation into parental sources.

## Introduction

Admixed genomes are characterized by a fragmented and recombined landscape of ancestral variation from their source populations, where chromosomes form mosaics shaped by demographic and evolutionary processes. Central to this pattern is meiotic recombination, which breaks and shuffles chromosomal segments inherited from distinct ancestral populations. At the core of this genomic restructuring lies the concept of *ancestry junctions*, which are points along the genome where segments of differing ancestral origins meet. The theoretical foundations for understanding junctions were first established by R.A. Fisher in the mid-20th century (Fisher, 1949).

These ancestry junctions arise during the early generations following admixture, when ancestry heterozygosity remains high and recombination between divergent ancestries is frequent. When two previously isolated populations interbreed, their offspring inherit large, contiguous chromosomal tracts from each ancestral source. In the first generation post-admixture, the offspring’s genomes consist of long, uninterrupted ancestry segments with few transitions. Over subsequent generations, recombination progressively fragments these segments. Each crossover event introduces new ancestry junctions, increasing the density of transitions along the genome.

Over longer timescales, the combined effects of recombination, demographic history, and evolutionary processes produce an intricate pattern of ancestry tracts. The spatial distribution of these tracts along the chromosome reflects both the time elapsed since admixture and the local recombination rate across the genome (Thompson, 2018). Analyses of the formation, frequency, and distribution of ancestry junctions therefore provide a powerful framework for inferring admixture timing, reconstructing demographic events, and characterizing recombination dynamics in natural populations.

Theoretical models have long treated ancestry junctions as genomic footprints shaped by recombination and demography. Early work in inbreeding and selfing established a basic condition: a crossover generates a new junction only when it occurs between homologous segments of different ancestry (Fisher, 1949, 1954; Bennett, 1953, 1954). Importantly, not all junctions are alike. In the classical two-haplotype view of a diploid, a junction is classified as *external* when the local ancestry state changes across it: on one side the diploid is ancestry-heterozygous, and on the other it is ancestry-homozygous, making the junction visible as an ancestry switch. Conversely, when the ancestry state does not change— either homozygous or heterozygous—the junction is *internal* and remains hidden in diploid organisms. Because segregation re-samples chromosomes across generations, junctions can shift categories. In other words, internal junctions may later become external, and external junctions may become internal (Fisher, 1954). This visibility framework reveals the long-term dynamics; whereby recombination progressively fragments ancestry blocks, causing junctions to accumulate, their spatial distribution approaching a Poisson process, and tract lengths converging toward an exponential distribution when the effective population size (*N*_*e*_) is large (Bennett, 1954; Fisher, 1954).

Building on this foundation, subsequent work extended the theory to specific mating schemes (Fisher, 1959; Gale, 1964) and to finite, randomly mating populations (Stam, 1980). Following Fisher’s distinction, analyses in this period concentrated on external junctions, the observable boundaries. Stam derived explicit formulae for the expected number of observable junctions per unit map length through time, linking junction accumulation to the probabilities that pairs and triplets of loci are not identical by descent (IBD) (Stam, 1980). Later models incorporated demographic factors such as changes in population size and subdivision, which alter the distribution of ancestral segment lengths (Chapman and Thompson, 2002). Complementary work derived the distribution of IBD tract lengths in finite random-mating populations, providing approximations for both the mean and variance (Chapman and Thompson, 2003). Another line of theory treated the survivability of ancestral blocks as a branching process under complete crossover interference and no coalescence. This framework yielded expressions for the number, size distribution, and total length of surviving blocks (Baird *et al*., 2003). Linking these expectations to observability, MacLeod and colleagues showed that the expected number of junctions per Morgan scales with the inbreeding coefficient *F*_*t*_, and that even dense marker maps may fail to detect a non-trivial fraction of junctions in inbred populations (Macleod *et al*., 2005). Finally, Palamara et al. (2012) introduced a practical framework for estimating admixture timing from the number of observed ancestry switches in diploid genomes. Their model, grounded in Wright–Fisher dynamics with uniform recombination, translated junction theory into an inference method. Collectively, these studies emphasize that junction formation and persistence are deeply shaped by demography and sampling, and that recombination signals carry complex genealogical information.

As genome-wide sequencing became more accessible, theoretical predictions about ancestry tracts and junctions could be directly tested using empirical data. Early work in this vein applied a hidden Markov model to reconstruct ancestry blocks in admixed individuals while accounting for background linkage disequilibrium in the ancestral sources (Tang *et al*., 2006). The approach was later refined to more explicitly link observed ancestry mosaics directly to the recombination and admixture processes that generate them (Gravel, 2012). This marked a turning point, where ancestry junctions transitioned from theoretical to practical signals for statistical inference. Subsequent studies demonstrated that tract length distributions encode demographic signals such as migration rates and timing (Pool and Nielsen, 2009). Researchers also clarified tract-length behavior under diverse histories, particularly when exponential assumptions fail, improving inference about the number and timing of admixture events (Liang and Nielsen, 2014). In parallel, theoretical work derived tract-length and junction-count distributions under a range of demographic conditions (Janzen *et al*., 2018), culminating in closed-form expressions for the expected number of junctions as a function of admixture time and recombination rate (Janzen and Miró Pina, 2022). More recently, these theoretical foundations have been combined with simulation and empirical data to evaluate robustness and broaden applicability across species (Hvala *et al*., 2018; Frayer and Payseur, 2021). Together, these advances linked population-genetic theory with tract-based inference, transforming ancestry mosaics from abstraction into practical signals for reconstructing demographic history.

Building on this progress, we introduce a new theoretical framework for ancestry switch dynamics that explicitly incorporates population-specific recombination maps, effective population size, and the decay of ancestry heterozygosity over time. Our derivations extend classical theory to predict the expected number of ancestry junctions under realistic genomic and demographic conditions. We evaluate these predictions using numerical and forward-time simulations that trace junction formation across generations. Finally, we compare our theoretical predictions to empirical switch counts in African American cohorts from multiple studies. By integrating fine-scale recombination landscapes with demographic processes, our work demonstrates that ancestry switches provide a rigorous and interpretable signal of admixture history. Furthermore, our method can be applied to any population as long as global ancestry fractions are available, which will allow for the inclusion of more diverse populations in genomics.

### Theory

We develop a theoretical framework to quantify the expected number of ancestry switches (Fisher junctions) produced after a 2-way admixture event. Specifically, we derive per-generation and cumulative expectations as functions of recombination and the decay of local ancestry heterozygosity.

#### Overview and scope

Throughout, we model junction counts along a segment of length |*L*| for a *single transmitted haplotype*. The underlying population follows a diploid Wright-Fisher model with effective size *N*_*e*_. We assume a single pulse of admixture followed by random mating within the admixed population; no subsequent gene flow from source populations; and neutral evolution with no selection acting on ancestry. We further assume that recombination in admixed individuals reflects ancestry-specific recombination processes that can be parameterized using population-specific recombination maps from parental sources. Recombination may be either uniform along the genome or allowed to vary, as specified in later sections.

All expectations are expressed per haplotype. Results for a diploid individual then follow (by symmetry) through doubling the haplotype counts. Population-level expectations are obtained by scaling according to the number of haplotypes in the sampled cohort. Physical distances are typically expressed in base pairs (bp) or megabases (Mb; 1 Mb = 10^6^ bp).

All parameter symbols used in this section are summarized in Table 1.

**Table 1:**
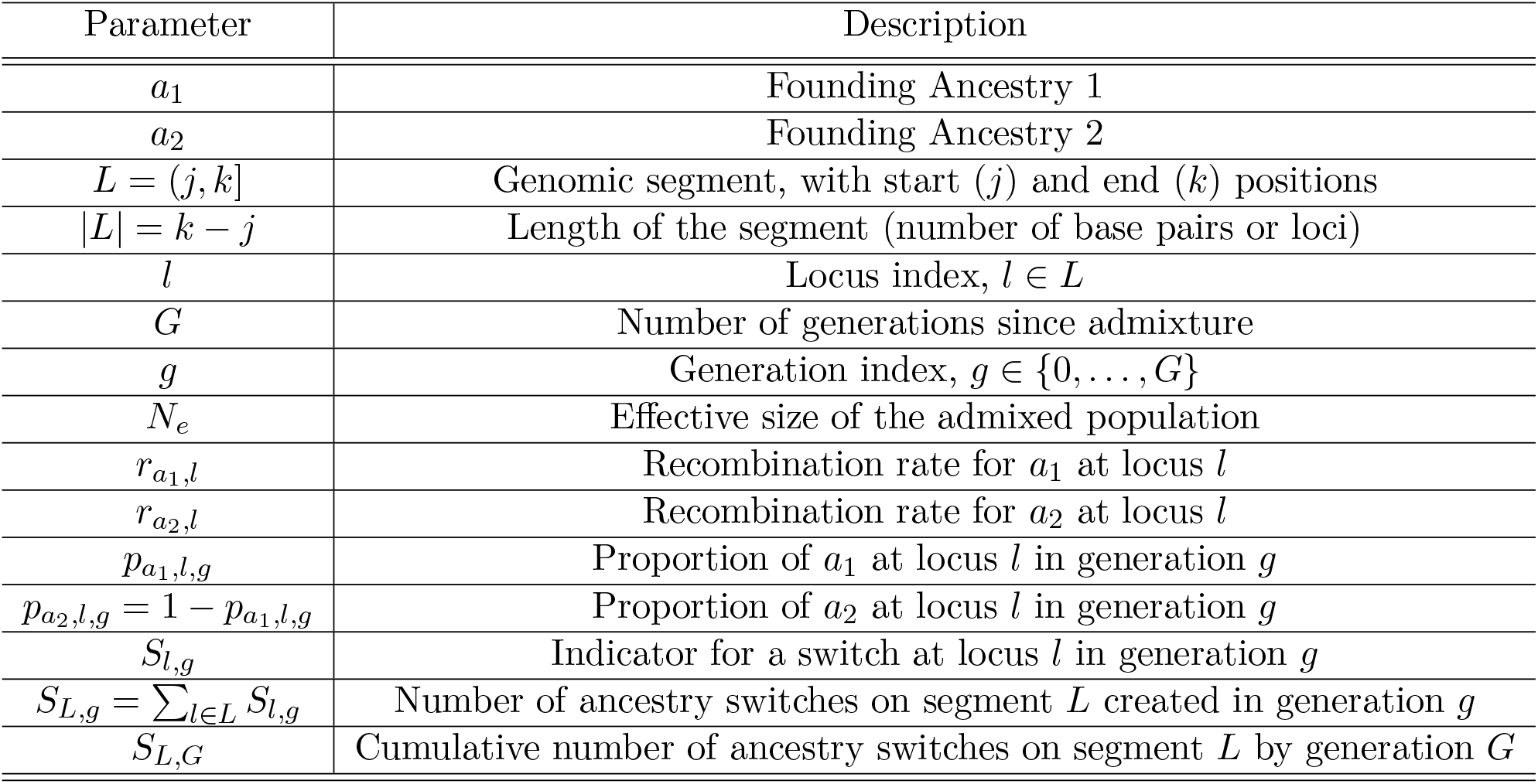
Model parameters used in the theoretical derivations.

### Per-generation expectation of ancestry switches

We begin by defining the probability ℙ[*S*_*l,g*_], where *S*_*l,g*_ is an indicator variable equal to 1 if an ancestry switch occurs at a single locus *l* in generation *g*, and 0 otherwise. Here, recombination at locus *l* refers to a crossover occurring anywhere within that interval. A detectable switch can only arise when the locus is ancestry-heterozygous; if both homologues carry the same ancestry, the probability of detecting a switch is zero. Using the law of total probability, we can decompose this probability into the chance of recombination given heterozygosity and the probability of heterozygosity itself:

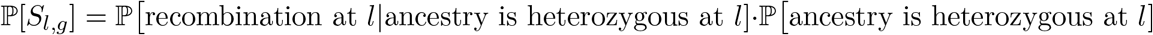

In a parent that is ancestry-heterozygous at *l*, the transmitted haplotype is equally likely to come from either the *a*_1_ or *a*_2_ homologue. Thus, the effective recombination rate at *l* is the mean of the ancestry-specific rates, 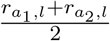. The probability that the two homologues carry different ancestries is 2 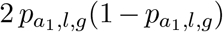, where the factor of 2 reflects the two ordered ancestry heterozygous configurations (*a*_1_|*a*_2_ or *a*_2_|*a*_1_). Substituting these into our decomposition gives:

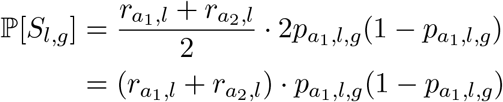

We now calculate 𝔼 [*S*_*L,g*_], the expected number of switches introduced in generation *g* along the segment *L* = (*j, k*], by summing the switch probabilities across all loci. This yields the expected number of switches per haplotype in a single generation, *g*.

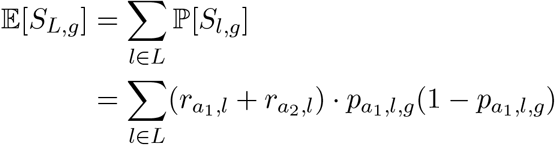

Under the assumption of uniform recombination along the segment (i.e., 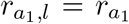 and 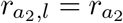 for all *l*), the factor 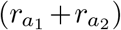 no longer depends on *l* and may be moved outside the summation:

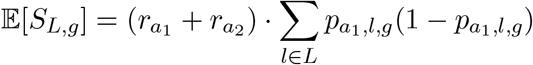

We can now express the sum as an expectation over all loci. For any function *f* (*l*) defined at each locus *l*, 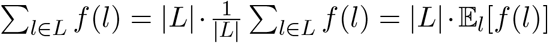, where |*L*| denotes the number of loci (or bases) in the segment. Applying this identity yields:

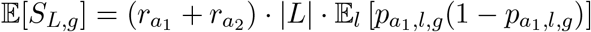

Next, we interpret the per-locus ancestry proportion 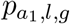 as the outcome of a random draw from locus *l*. Formally, let 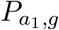 be the random variable equal to 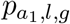 when *l* is chosen uniformly at random from the segment. Then, the average over loci is the expectation of 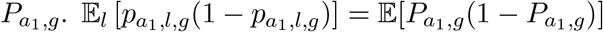.

Using the identity 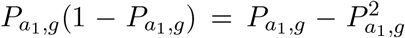 and the linearity of expectation, we obtain:

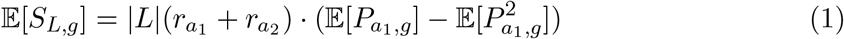

Equation (1) shows that the expected number of ancestry switches in generation *g* is proportional to the average recombination rate across ancestries and to the level of ancestry heterozygosity present in a given generation. This connects back to the classical result that recombination generates new junctions only when different ancestries co-occur (Fisher, 1949). In the next section, we derive the dynamics of these expectations under genetic drift, which governs the decay of heterozygosity through time.

### Temporal dynamics of expected ancestry switches

To convert our per-generation switch expectation (Equation (1)) into a fully time–dependent prediction, we must model how local ancestry heterozygosity evolves post-admixture. Under a neutral Wright-Fisher model with the effective size of the admixed population *N*_*e*_, the mean ancestry proportion remains fixed at the initial value 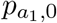 and ancestry heterozygosity decays by a factor 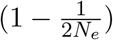 each generation (Hartl and Clark, 2007; Ewens, 2004). Therefore, we can write the expected ancestry heterozygosity at generation *g* as:

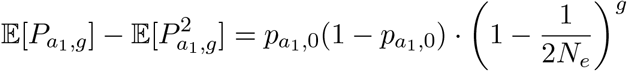

If we substitute the decay of ancestry heterozygosity into Equation (1), we obtain:

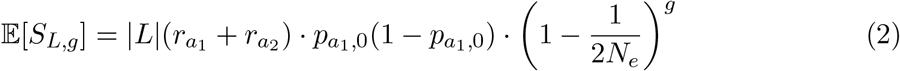

Equation (2) gives the expected number of new switches created in generation *g* on a single haplotype. Ancestry switches are generated at the highest rate immediately after admixture and then decline in proportion to the erosion of local ancestry heterozygosity. This expression illustrates how recombination, initial ancestry composition, and effective population size jointly shape the temporal pattern of switch formation. For an in-depth derivation, refer to *Supplemental Theory*.

In the next subsection, we integrate these per-generation contributions to obtain the cumulative number of ancestry switches over *G* generations.

### Cumulative expected ancestry switches over *G* generations

To extend Equation (2) beyond a single generation, we sum the expected switches over *G* generations.

For notational convenience, let 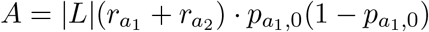.

Then:

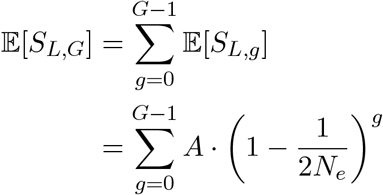

Note, this is a geometric series 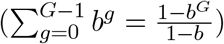 with 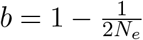. Hence,

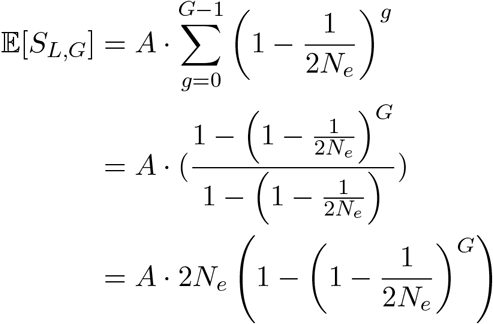

Substituting *A* back in, we get:

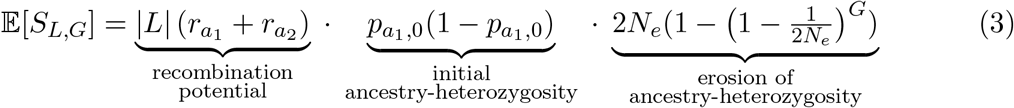

Given the assumptions of neutrality and a single-pulse of admixture, Equation (3) gives the total expected number of switches accumulated up to generation *G*, capturing the combined influence of recombination potential, initial ancestry heterozygosity, and its erosion through genetic drift.

Closely related expectations for the accumulation of ancestry junctions through time were previously derived under Wright–Fisher dynamics assuming uniform recombination, with results expressed in terms of total genetic map length (Janzen *et al*., 2018; Janzen and Miró Pina, 2022). Under these simplifying assumptions, Equation (3) recovers these earlier results when the total recombination potential is written as a single map-length parameter, with the initial ancestry heterozygosity 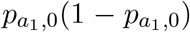 playing the same role as the heterozygosity term used in Equation 3 of Janzen *et al*., 2018. The present formulation makes this dependence explicit in terms of physical distance and ancestry-specific recombination rates, which enables expectations to be evaluated by integrating recombination potential across the entire genome and extended naturally to heterogeneous recombination landscapes through direct integration with empirical recombination maps.

### Extending the equation for non-constant recombination

Across the genome, recombination landscapes are highly heterogeneous, with localized hotspots of elevated crossover activity interspersed among broad coldspots of reduced rates. These fine-scale differences are captured by population-specific recombination maps, which quantify how the probability of crossover varies along each chromosome. Incorporating such maps allows our model to account for spatial heterogeneity in recombination while keeping the ancestry proportion fixed across the genome.

To accommodate this spatial heterogeneity, we generalize the uniform-rate term 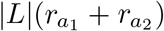 from Equation (3) to a position-dependent formulation across the segment *L* = (*j, k*] of length |*L*| = *k* −*j*. In what follows, we present: (i) a continuous formulation over the physical coordinate *x* ∈ *L*, and (ii) discrete formulations based on genomic breakpoints (interval-wise approximation) or fine-scale per-base summation when rates are available.

#### Continuous evaluation

Let *x* ∈ *L* = (*j, k*] denote a physical base-pair position within the segment of length |*L*| = *k* − *j*. We allow the recombination rates 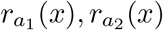 to vary with *x*. Then:

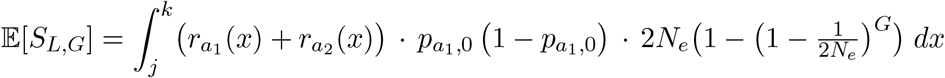

We factor out terms that do not depend on *x*, which yields:

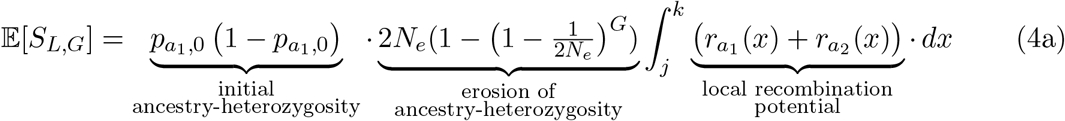

Here, 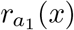 and 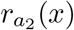 denote the recombination rates in Morgans per base pair (bp) at position *x*, when the transmitted haplotype is carried on ancestry *a*_1_ or *a*_2_, respectively.

#### Discrete evaluation

A typical recombination map lists the breakpoint (recombination event) positions together with an associated crossover rate. These positions mark loci where the recombination rate is defined or estimated, typically at single-nucleotide polymorphisms (SNPs) or other informative markers. The recombination landscape can be inferred or estimated from different types of population, pedigree, or gamete-based data (Peñalba and Wolf, 2020). The recombination rates between breakpoints are obtained by mathematical interpolation to yield a fine-scale rate profile. To apply Equation (4a) in practice, we discretize the segment as follows.

Let *j* = *x*_1_ < *x*_2_ < · · · < *x*_*W* +1_ = *k* denote breakpoint positions. For each *w* = 1, …, *W*, define the interval [*x*_*w*_, *x*_*w*+1_) of length Δ*x*_*w*_ = *x*_*w*+1_ − *x*_*w*_ (bp). We choose a representative position 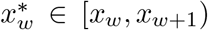 at which to evaluate the per-base recombination rates. Then:

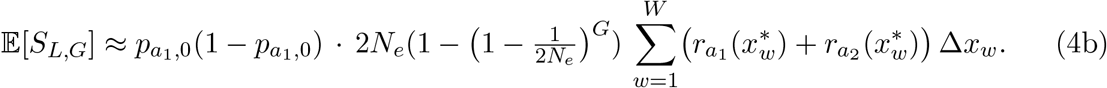

This is exact if 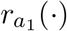 and 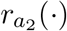 are constant on each interval [*x*_*w*_, *x*_*w*+1_).

If fine-scale per-base data (either directly or via interpolation) is available one can sum over all positions:

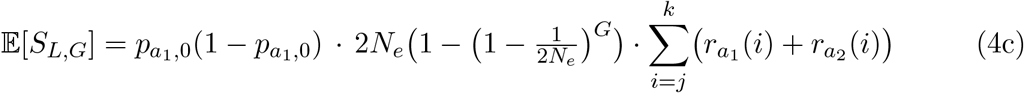

where *i* indexes base-pair positions in the segment (*j, k*], rates are in Morgans/bp, and the per-base step size is Δ*x* = 1 bp (hence omitted).

*Note:*

i. If recombination rate does not differ by ancestry, replace 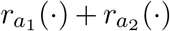 with 2 *r*(·) in (4b) and (4c).
ii. If per-base rates are provided as cM/Mb, convert to Morgans/bp via 1 cM/Mb = 10^−8^ Morgans/bp before using Equations (4b) or (4c). 1 cM corresponds to a 1% probability of crossover between two loci per generation.
iii. Under bounded, piecewise-continuous 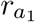 and 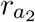, the interval sum in (4b) converges to (4a) as the maximum interval length max_*w*_(*x*_*w*+1_ − *x*_*w*_) → 0; (4c) is the per-base Riemann sum.

## Methods

### Theoretical framework and ancestry switch estimation

#### Parameter dynamics

To investigate how the expected number of ancestry switches accumulates over time following admixture, we evaluated Equation (3) under a range of parameter values. Equation (3) predicts the expected number of ancestry junctions per haplotype as a function of generations since admixture (*G*), initial ancestry proportion 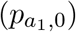, recombination rate (*r*), and effective population size (*N*_*e*_). Our theoretical model assumes Wright–Fisher dynamics with constant population size, a single-pulse admixture at *G* = 0, no subsequent gene flow, and neutral evolution. Recombination is assumed to occur uniformly along the chromosome, and ancestry switches are counted per haplotype. These assumptions define the baseline conditions under which theoretical expectations for ancestry switch accumulation are derived and evaluated.

We use a deterministic implementation in Python with NumPy (Harris *et al*., 2020) to compute values of Equation (3) across multiple generations. To illustrate how each parameter modulates switch accumulation, we vary the initial ancestry proportion 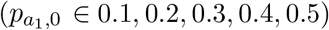; the recombination rate (*r* ∈ 10^−9^, 10^−8^, 10^−7^, 10^−6^, 10^−5^); and the effective population size (*N*_*e*_ ∈ 10^2^, 10^3^, 10^4^, 10^5^, 10^6^). We evaluated the expectation at *G* ∈ 0, 5, 10, …, 50 generations.

Default parameters were: chromosome length *L* = 2.5×10^8^ bp (approximating human chromosome 1); recombination rates 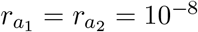 per base-pair and per generation; initial ancestry proportion 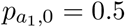 and effective population size *N*_*e*_ = 10, 000 (approximating global human effective population size).

#### Quantifying the number of expected ancestry switches from theory

For each generation, we computed the expected number of ancestry switches per haplotype under the two previously introduced models. For the numerical simulations, we use the following parameters: chromosome length *L* = 2.5 × 10^8^ bp (approximating human chromosome 1 (GRCh38)); initial ancestry proportion 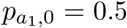 and an effective population size of *N*_*e*_ = 14, 000. This value is consistent with long-term estimates of human effective population size, which are on the order of approximately 14,000 individuals (Li and Durbin, 2011; Gravel et al., 2011; Medina-Muñoz et al., 2023). Then, we evaluate both a constant and non-constant recombination map.

For the constant recombination model, we evaluate Equation (3), and assume a uniform recombination rate of 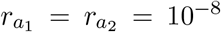 across the segment. For the non-constant recombination map model, we apply the discretized formula in Equation (4b), and assume recombination rates vary across the segment. We generate a custom recombination map by partitioning the chromosome into 1000 randomly sized intervals and assigning each a recombination rate drawn from a uniform distribution between 1 × 10^−9^ and 1 × 10^−7^ Morgans per base pair, corresponding to realistic recombination rate ranges. This choice ensures even coverage across a plausible range of recombination rates; alternative sampling schemes (e.g., log-uniform) are also possible and would yield similar trajectories, reflecting the linear dependence of switch accumulation on recombination rate. In the SLiM simulations, this approach was chosen to isolate the effects of recombination rate variation under controlled conditions. For comparisons against human empirical data, population-specific recombination maps are used in the analysis.

Expected switch counts are tabulated deterministically by summing across map intervals, with contributions weighted by the local recombination rate and ancestry heterozygosity. All theoretical computations are implemented in Python using NumPy (Harris *et al*., 2020) for numerical operations and pandas (McKinney, 2010) for data manipulation.

### Simulation framework and ancestry switch estimation

We simulate ancestry switch dynamics in an admixed population using SLiM 3 (Haller and Messer, 2019), a forward-time simulator capable of recording full tree sequences across the genome (Haller *et al*., 2019). Each simulation began with a symmetric split of a common ancestral population into two source populations at generation 1000. These source populations, denoted *a*_1_ and *a*_2_, contributed equally to a newly formed admixed population via a single-pulse admixture event 10 generations before the present (generation 1900). We then allow the admixed population to evolve forward under Wright-Fisher dynamics with constant population size and random mating, recording tree sequences every generation from 1900 to 1910. These generation times were chosen to capture the early-phase accumulation of ancestry junctions following admixture, while leaving room for future extensions that incorporate more complex demographic scenarios that may require a burn-in period.

To evaluate how recombination shapes the accumulation of ancestry switches, we implement a similar simulation framework as above. Briefly, all simulations use the following baseline parameters: chromosome length *L* = 2.5 × 10^8^ bp (approximating human chromosome 1); recombination rates; initial ancestry proportion 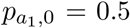 and effective population size *N*_*e*_ = 14, 000 (approximating human effective population size (Li and Durbin, 2011; Gravel *et al*., 2011; Medina-Muñoz et al., 2023)). Recombination rates are assigned as described above for each model. The first uses a constant recombination model, where a uniform recombination rate of 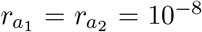 was applied to the segment. The second uses the recombination map model, where recombination rates vary across the segment.

### Quantifying the number of observed ancestry switches from simulations

To quantify the number of ancestry switches introduced by recombination, we analyze the tree sequences output by SLiM at each generation post-admixture. The tree files contain the complete set of marginal genealogies along the chromosome, enabling us to trace the ancestry of sampled haplotypes back to the source population of each founding individual. We define an ancestry switch as a change in founder ancestry between adjacent genomic intervals. That is, if one marginal tree traces a genomic segment to source population *a*_1_, and the next marginal tree (immediately to its right) traces to source population *a*_2_, this is counted as a switch.

To determine founder ancestry, we traverse each marginal tree upward from a sampled segment until we reach the first remembered ancestor. These ancestors, sampled at generation 1899 (just before admixture), were tagged using SLiM’s `treeSeqRememberIndividuals()`, which preserves their identity and metadata across generations(Haller *et al*., 2019). The segment is assigned the ancestry of the remembered ancestor. We then identify ancestry switches by comparing the labels of adjacent genomic intervals.

The procedure is repeated for *n* = 10 independent simulation replicates. For each replicate, we compute the number of ancestry switches for every sampled haplotype and then calculate the mean switch count per haplotype across the population. We average the replicate-level means to obtain the final observed expectation of switches per haplotype at each generation. A detailed description of the switch count algorithm is in Algorithm S1.

### Application to an admixed population

Here, we apply our theoretical framework to empirical data from an admixed human population to illustrate how ancestry switch expectations behave under realistic demographic and recombination scenarios. Using parameter values drawn from existing studies, we compare theoretical switch counts to those observed in human genomes in order to assess the consistency of the model with empirical patterns. This application is intended to demonstrate how the framework can be parameterized and interpreted in practice when applied to real data.

#### Calling ancestry with FLARE

Whole genome sequence (WGS) data from the 1000 Genomes Project (GRCh38) (Byrska-Bishop *et al*., 2022) was downloaded and filtered to autosomal bi-allelic markers only. Then, we use FLARE v0.4.2 (Browning *et al*., 2023) to conduct local ancestry inference for 74 admixed African Americans from the Southwest (ASW). The reference panel contains 178 Yoruba in Ibadan, Nigeria (YRI) and 179 Utah residents (CEPH) with Northern and Western European ancestry (CEU) individuals which are used as representative populations for African and European/European-American ancestry, respectively. We use the default parameters of FLARE alongside of their genetic map for human GRCh38. After running FLARE, we remove samples that were previously noted to have high autozygosity (Gazal *et al*., 2015). Then, we computed relatedness using KING (Manichaikul *et al*., 2010), and extract unrelated individuals using plink2 v2.00a4.3 (Chang *et al*., 2015) with a KING kinship coefficient cutoff of 0.0448.

We then conduct post-hoc filtering of the inferred local ancestry. First, we reduce the dataset to only chromosome 1, which is our representative chromosome for simulations. Markers from the FLARE inference results are removed if they exist outside the 1000 Genomes strict mask or inside the ENCODE Exclusion List Regions for GRCh38 (Amemiya *et al*., 2019). We then combine adjacent markers of the same inferred ancestry into non-overlapping, contiguous ancestry segments for each sample. Segments shorter than 2 Mb were removed. Subsequently, we remove samples if, on either sample haplotype, the total length of remaining segments is less than 2 standard deviations of the population mean. Post-filtering, adjacent segments of the same inferred ancestry are annealed together, producing the final set of non-overlapping, contiguous ancestry segments for each remaining sample.

#### Empirical switch count for ASW

After filtering, we anneal FLARE’s positional annotation output to provide non-overlapping, contiguous ancestry segments for each sample’s haplotypes. The total switch counts for each sample’s haplotype is equal to the total number of these ancestry segments on that sample’s haplotype minus 1. The total switch count is the sum across all samples’ haplotypes in the population. The total switch count is then divided by the number of chromosomes to produce the observed switch count mean. In addition, we also calculate a bootstrapped resampling mean of 1000 iterations and bias-corrected and accelerated (BCa) 95% bootstrap interval.

#### Theoretical switch count for ASW

To incorporate realistic recombination landscapes into our theoretical predictions, we use population-specific recombination maps inferred with *pyrho* (Spence and Song, 2019) for the YRI and CEU populations. Each map consist of an array of genomic positions with corresponding recombination rates (which we converted to Morgans per base pair). Mirroring the simulation setup, we use chromosome 1 (length = 248,956,422 bp) as a proxy for calculating expected switch counts in the ASW populations.

To account for heterogeneity in recombination along the chromosome, we partition chromosome 1 into intervals of varying lengths. Intervals are defined by taking the union of all positions in the CEU and YRI maps, with each position marking the start of a new interval as described in Equation (4b). This procedure yields 202,540 intervals. Within each population, recombination rates were forward-filled so that every interval is assigned a value, with missing entries set to zero at the chromosome start and end.

The merged table for chromosome 1 includes the genomic start position of each interval, CEU- and YRI-specific recombination rates (*r*_*Y RI*_ (*x*) and *r*_*CEU*_ (*x*)), and the initial YRI ancestry proportion (*p*_*Y RI*,0_). Expected ancestry switch counts were calculated across intervals according to Equation (4b), with interval lengths Δ*x* given by the distance between consecutive start positions.

For empirical parameterization, we set the effective population size to *N*_*e*_ = 728, corresponding to the combined contributions of African, European, and African-American genealogical ancestors reported in Table 3 of (Agranat-Tamir *et al*., 2024). Here, *N*_*e*_ denotes the effective population size of the admixed population, which determines the strength of genetic drift and the resulting decay of ancestry heterozygosity. The number of generations since admixture was fixed at *G* = 14, following estimates from (Mooney *et al*., 2023). Initial ancestry proportions were assumed to be uniform across the chromosome, with values chosen to reflect both the upper-bound estimate (*p*_*Y RI*,0_ = 0.85) and lower-bound estimate *p*_*Y RI*,0_ = 0.75, consistent with previous studies of African American populations (Smith *et al*., 2004; Cheng et al., 2009; Baharian et al., 2016; Mooney et al., 2023).

#### Estimating switch counts from other African American cohorts

To facilitate approximate comparison with previously published results, we approximated chromosome-specific ancestry switch counts from earlier studies based on their reported local ancestry assignments and summary statistics. In Baharian et al. (2016), Figure 1D displays inferred local ancestry along both haplotypes of chromosome 1, with one haplotype exhibiting four ancestry switches and the other exhibiting eight switches. We therefore estimated the per-individual switch count for chromosome 1 as the average across haplotypes, yielding six switches (Baharian *et al*., 2016). Similarly, in Gravel (2012), Figure 1 presents local ancestry assignments for chromosome 1, with five switches observed on one haplotype and two on the other, resulting in an average of three switches per individual (Gravel, 2012).

**Figure 1.**
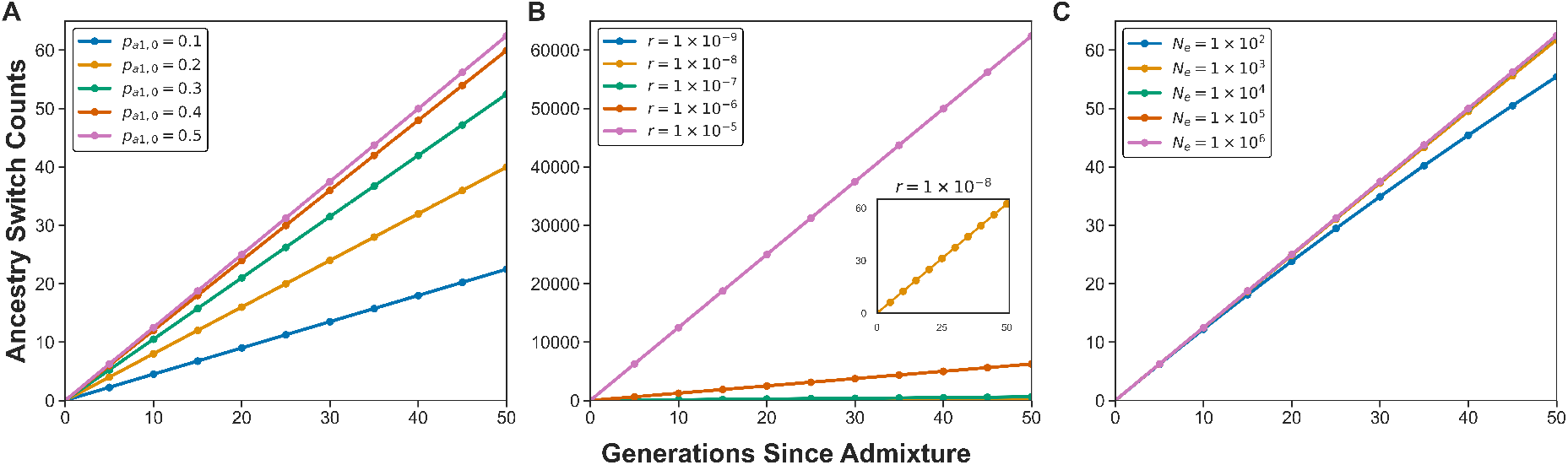
Cumulative theoretical ancestry switch counts after *G* generations under varying model parameters. For panel A) we vary ancestry proportion 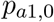, B) we vary recombination rate *r*, and C) we vary effective population size *N*_*e*_. The total expected new switch counts are calculated from Equation (3) for a haploid chromosome of length *L* = 2.5 × 10^8^ bp. Baseline parameters 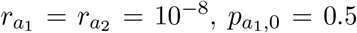, and *N*_*e*_ = 10; values shown for *G* ∈ {0, 5, …, 50}. All expectations are per haplotype.

Wegmann et al. (2011), who used ancestry switch points in admixed individuals to infer high-resolution recombination maps, reported only genome-wide switch counts. In Supplementary Note 2.5, they reported an average of 91.98 ancestry switch points per individual across approximately 2.5 × 10^9^ bp after excluding unsequenced regions and centromeres. Dividing the reported number of switches by the analyzed genome length yields an average density of approximately 0.368 switches per megabase (Mb). Because the effective length of chromosome 1 used in their analysis was not reported explicitly, we estimated it using Supplementary Figure S10, which shows the derived recombination map for chromosome 1 with excluded regions indicated by gray rectangles. Based on this figure, we estimated that approximately 40–50 Mb were excluded, corresponding to an analyzed chromosome 1 length of roughly 200–210 Mb. Multiplying this estimated length range by the genome-wide switch density yields an expected range of approximately 7.36–7.73 switches on chromosome 1. In Figure 4, we report the midpoint of this range, 7.545 switches, as a point estimate (Wegmann *et al*., 2011). We emphasize that these values represent approximations derived from published summaries and graphical information rather than directly reported chromosome-specific estimates.

## Results

### Parameter dynamics

To establish baseline expectations for ancestry switch accumulation under different model parameters, we used Equation (3) to calculate the expected number of switches per haplotype across time. Figure 1 shows these trajectories and how they vary with ancestry proportion, recombination rate, and population size. All values were calculated per haplotype; under symmetric assumptions, diploid values would be approximately twice as large.

First, we varied the initial ancestry proportion 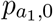 while holding other parameters constant (Figure 1A). Switch accumulation was fastest when ancestry was evenly balanced between source populations, with the steepest slope at 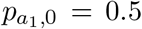. As 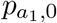 decreased, we observed a corresponding reduction in the trajectories of the slope, where the changes become shallower, reflecting the reduced ancestry heterozygosity. The trajectories were symmetric and ordered according to 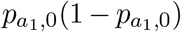, consistent with the expected relationship between heterozygosity and switch formation.

Next, we varied the recombination rate while holding other parameters constant (Figure 1B). Switch counts scaled linearly with *r*, with each ten-fold increase producing a proportional increase in the slope of the trajectory. The inset highlights the trajectory corresponding to the canonical human baseline recombination rate of *r* = 10^−8^ Morgans per base pair per generation.

Lastly, we varied the effective population size while holding other parameters constant (In Figure 1C). Larger populations retained more ancestry heterozygosity over time, leading to slightly higher switch counts compared with smaller populations. Although trajectories remained close together in the early generations, a long-term plateau in switch accumulation is expected to depend on the strength of drift. The per-generation ancestry switch count across these parameters is available in Figure S2.

### Simulation framework and ancestry switch estimation

Next, we used forward-time simulations to test whether theoretical predictions matched observed ancestry switch dynamics. We simulated a symmetric admixture event followed by random mating, and tracked haplotypes over the first 10 generations after admixture (Figure 2). The tree sequence preserved complete genealogical information, which enabled direct counts of ancestry switches along each chromosome under both constant and heterogeneous recombination models.

**Figure 2.**
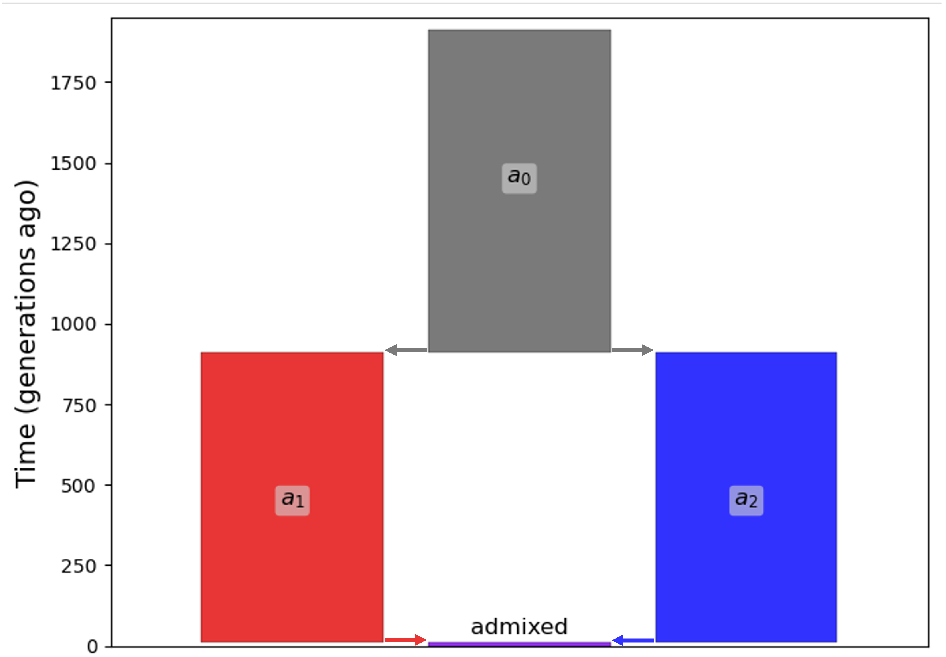
Schematic of the SLiM simulation framework over a time period of 1910 generations. The y-axis represents the time since the present. The ancestral population (*a*_0_) splits symmetrically 910 generations ago into two source populations, populations 1 (*a*_1_) and 2 (*a*_2_). After some time, *a*1 and *a*2 contribute equally (rate = 0.5) to an admixed population 10 generations ago via a single-pulse admixture event. Tree sequences are recorded every generation starting at 10 generations ago to the present to capture early ancestry switch dynamics.

Using this two-way admixture model (Figure 2), we calculated ancestry switch counts from both forward-time simulations and theoretical predictions derived from Equation (3). Under the constant recombination model (Figure 3A), the observed simulated switch trajectories were nearly indistinguishable from the theoretical expectation. Mean switch counts across replicates closely followed the analytical expectation, and the 95% confidence intervals were narrow, indicating minimal variability among simulation replicates.

**Figure 3.**
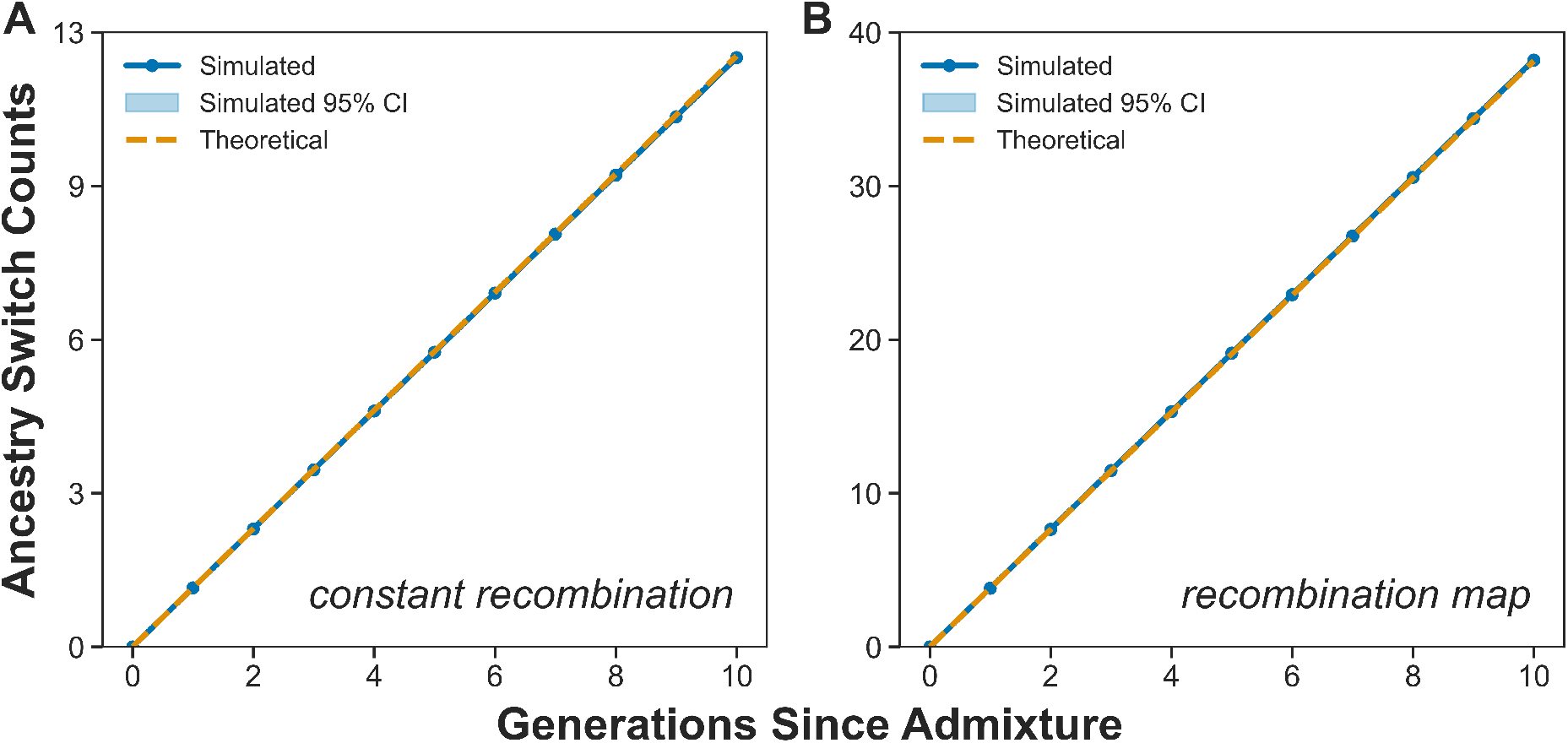
Comparison of simulated and theoretical ancestry switch counts under constant and variable recombination scenarios. A) Switch accumulation across generations assuming a constant recombination rate across the genome. B) Switch accumulation assuming variable recombination, informed by a generated recombination map. Theoretical expectations were calculated using Equation (3) for the constant recombination scenario and Equation (4b) for the variable recombination scenario. The simulated values were calculated via TreeSeq outputs from SLiM simulations.

We obtained similar results with the recombination map model (Figure 3 B), where recombination rates varied along the chromosome. Despite substantial local variation in rates, the cumulative number of ancestry switches across the genome aligned closely with theoretical predictions from the discretized formulation in Equation (4b). These results demonstrate that the theoretical framework provides accurate predictions of ancestry switch accumulation under both uniform and spatially variable recombination landscapes.

Switch count trajectories across a range of population sizes are shown for the constant recombination model (Figure S3) and for the non-constant recombination map model (Figure S4)

### Application to African-American populations

Next, we compared the observed number of ancestry switches in African Americans from the 1000 genomes project (Byrska-Bishop *et al*., 2022) to theoretical expectations calculated from Equation (4b). Figure 4 presents results for chromosome 1, where theoretical switch counts were parameterized with an effective size (*N*_*e*_ = 728) and demography (*G* = 14) from previously published work on genealogical ancestors of African Americans (Mooney *et al*., 2023; Agranat-Tamir *et al*., 2024). While Figure 4 focuses on chromosome 1 for clarity, corresponding empirical and theoretical switch counts for all autosomes are provided in Table S1. To account for uncertainty in the initial ancestry composition, we considered two realistic values for the African ancestry proportion, with YRI used as a proxy: a lower-bound value of *p*_*Y RI*,0_ = 0.75 and an upper-bound value of *p*_*Y RI*,0_ = 0.85 (Cheng *et al*., 2009; Baharian et al., 2016).

**Figure 4.**
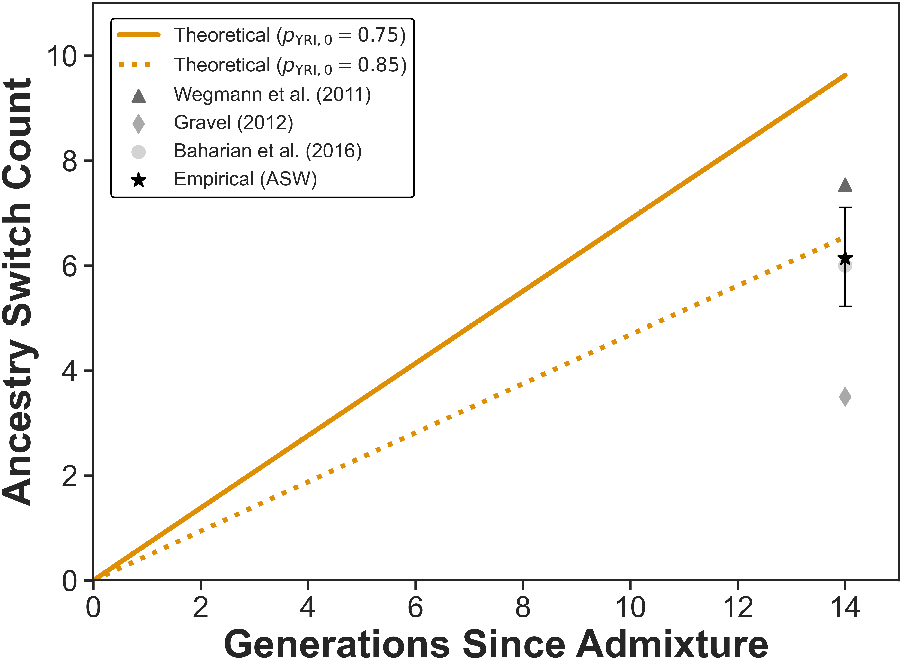
Comparison of empirical and theoretical ancestry switch counts for African American populations. Theoretical ancestry switch counts (orange lines) were calculated under two initial YRI ancestry proportions: *p*_*Y RI*,0_ = 0.75 (solid) and *p*_*Y RI*,0_ = 0.85 (dotted), assuming *N*_*e*_ = 728 and *G* = 14. The black star shows the empirical mean switch count estimated from chromosome 1 in ASW individuals from the 1000 Genomes Project, with vertical bars indicating the 95% bias-corrected and accelerated bootstrap confidence interval. For reference, published estimates from single African-American samples (GeneSTAR, GENOA, GRAAD21–23, and SARP and CAG/CSGA cohorts (Wegmann *et al*., 2011), dark grey triangle; NA19700 (Gravel, 2012), medium grey diamond; HRS cohort (Baharian *et al*., 2016), light grey circle) are also shown.

The empirical switch counts observed in the ASW population (Byrska-Bishop *et al*., 2022) (black star) were more consistent with the upper-bound of ancestry from the theoretical model (*p*_*Y RI*,0_ = 0.85), falling below the expectations derived from the lower-bound of ancestry (*p*_*Y RI*,0_ = 0.75). The 95% bootstrap confidence interval around the ASW estimate overlapped the *p*_*Y RI*,0_ = 0.85 curve, but not the *p*_*Y RI*,0_ = 0.75 curve, indicating that the higher initial African ancestry proportion provides a better fit to the observed data. When interpreting these comparisons, it is important to note that empirical and theoretical switch counts are computed over different effective genomic lengths. Theoretical expectations use full chromosome lengths from the GRCh38.p14 assembly, whereas empirical estimates are restricted to callable genomic regions retained after quality control. Because expected switch counts scale linearly with segment length, this distinction contributes to systematic differences between empirical and theoretical values across chromosomes.

Further, we compared the ASW switch counts to previously reported estimates from other African American cohorts. Wegmann et al. (2011) reported ancestry switch counts for individuals drawn from several independent studies, including GeneSTAR, GENOA, GRAAD21–23, and SARP and CAG/CSGA. The switch count reported by Wegmann et al. (2011) (dark gray triangle) was slightly higher than the ASW estimate, but remained well below the theoretical expectation derived under the lower-bound ancestry proportion (*p*_*Y RI*,0_ = 0.75) (Wegmann *et al*., 2011). In contrast, the switch count for individual NA19700 (Gravel, 2012) (medium gray diamond) was markedly lower than both our ASW estimate and theoretical expectations. Conversely, the value from an individual in the HRS cohort (Baharian *et al*., 2016) (light gray circle) was nearly identical to the ASW mean. Together, our results show that switch counts in African American populations are broddly consistent with theoretical predictions, despite expected variation across individuals and cohorts (Baharian *et al*., 2016).

## Discussion

Ancestry switches provide a direct record of how recombination reshapes genomes after admixture. Here we show that their accumulation can be predicted with precision from first principles, linking switch dynamics to recombination, ancestry heterozygosity, and effective population size. By extending classical junction theory to incorporate population-specific recombination maps, our framework moves beyond uniform models and captures fine-scale heterogeneity in crossover activity. The model provides a parameterized framework that uses measurable quantities (e.g. global ancestry and recombination rate) to offer information about complex admixture histories through a mechanistic model that builds upon prior admixture models (Tang *et al*., 2006; Gravel, 2012; Browning et al., 2023). Further, the model does not require partitioning the admixed populations genome into portions that can be traced to each source. Instead, the model focuses on the admixed population by integrating information from each source population.

One of the more interesting patterns that emerges from the model appears when comparing parameter effects across timescales (Figure 1). In the short term, recombination rate and ancestry proportion dominate switch accumulation, producing trajectories that are approximately linear; and the role of effective population size is not apparent. However, after many generations the impact of effective population size (Figure 1C) becomes evident, because drift has progressively eroded ancestry heterozygosity. After approximately 100 generations, we observe that the generation of new switches in small populations begin to plateau early, whereas large populations continue accumulating switches for hundreds of generations (Figure S1).

Next, we used forward-in-time simulations to corroborate the theoretical predictions. We observed that simulated switch counts closely matched analytical expectations under both constant and variable recombination scenarios (Figure 3). Confidence intervals were generally narrow, reflecting the predictability of switch accumulation once recombination rates and ancestry heterozygosity are specified. Larger deviations were limited to very small populations, where genetic drift amplifies variation among replicates (Supplementary Figures S3 and S4). As shown in *Supplemental Theory*, drift increases the variance in local ancestry proportions by roughly 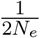 per generation, reducing the predictability of switch counts as heterozygosity decays.

Importantly, a key source of realism in our framework is the explicit incorporation of population-specific recombination maps, rather than assuming constant recombination rates across populations and/or the genome. Accounting for these maps improved the accuracy and stability of switch-based inference, particularly in small populations where stochastic effects are strongest (Supplementary Figures S3 and S4). We suspect this is due to recombination maps containing information about hotspots, which may constrain where switches can form, as these genomic regions have high crossover potential. Thus, the recombination map reduces stochastic variation across replicates and yields tighter confidence intervals than those under uniform recombination models.

Empirical studies have shown that recombination landscapes can differ across populations, reflecting variation in hotspot activity and recombination modifiers such as PRDM9 (Baudat *et al*., 2010; Parvanov *et al*., 2010; Hinch *et al*., 2011; Spence and Song, 2019; Hassan *et al*., 2021). Similar divergence has been observed in other taxa, including mouse subspecies (Dumont *et al*., 2011) and *Drosophila* (Chan *et al*., 2012), underscoring the evolutionary plasticity of recombination control mechanisms. These differences are often concentrated and can persist across many generations. At broader resolutions (i.e., chromosome-scale and megabase-scale windows) recombination rates tend to be highly correlated across populations, indicating substantial conservation of overall recombination patterns within species (Hinch *et al*., 2011; Singhal et al., 2015).

Previous work shows that the extent to which recombination patterns in parental populations are informative is context-, scale-, and species-dependent(Singhal *et al*., 2015; Schumer *et al*., 2018; Samuk et al., 2020). In admixed human populations, recombination may differ locally from that of either parental population due to composite ancestry histories and fine-scale differences in hotspot usage, while remaining broadly similar to parental recombination patterns at genome-wide scales (Hinch *et al*., 2011; Dinh et al., 2024).

In humans specifically, relatively low genetic divergence among populations, shared PRDM9-mediated recombination mechanisms, and the use of sex- and time-averaged recombination maps imply that parental recombination maps are informative at the broad, genome-wide scales relevant to ancestry switch accumulation Hinch et al. (2011). Because ancestry switch counts integrate recombination over extended genomic regions and short post-admixture timescales, they are robust to fine-scale differences in hotspot location. This scale dependence explains why switch-based inference can remain accurate even when recombination landscapes in admixed individuals differ locally from those of either parental population.

Despite biological complexity, most tract- and junction-based theories express expectations in genetic map units and assume a shared or uniform map for inference (Stam, 1980; Chapman and Thompson, 2002; Gravel, 2012; Liang and Nielsen, 2014; Janzen et al., 2018; Janzen and Miró Pina, 2022). In particular, the junction-count framework developed by Janzen et al. (2018) and extended by Janzen and Miró Pina (2022) derives expectations for ancestry switch accumulation as a function of total genetic map length under uniform recombination. Our framework recovers these results under the same assumptions, while generalizing them by expressing recombination potential explicitly as a function of physical distance and ancestry-specific recombination rates, enabling direct integration over heterogeneous recombination landscapes across the genome. These findings, together with our framework that explicitly accounts for recombination rate variation, underscore the value of including population-specific recombination maps. Further, the model provides a conceptual bridge linking recombination biology and ancestry inference, clarifying how fine-scale recombination differences among ancestral source populations translate into accurate theoretical expectations of switch counts in admixed populations.

After testing the model with simulations, we next applied it to empirical data. We chose to predict switch counts in African Americans from the publicly available 1000 Genomes dataset(Byrska-Bishop *et al*., 2022). In this application, the human data are used to illustrate how theoretical switch count expectations parameterized using values drawn from the literature, align with empirical patterns observed in an admixed population. When we applied the model we found that the predicted switch counts overlapped the confidence intervals of the observed switch counts within unrelated African-American individuals (Figure 4). In this analysis, theoretical expectations are computed using full chromosome lengths, whereas empirical switch counts are based on the subset of genomic regions retained after ancestry inference quality control. Because expected switch counts scale linearly with segment length, this difference can lead to modest downward shifts in empirical switch counts relative to theoretical predictions, particularly for smaller chromosomes (Table S1). The theoretical model was parameterized with a realistic demography and admixture timing (Mooney *et al*., 2023; Agranat-Tamir *et al*., 2024), and aligned more closely with the higher initial African ancestry proportions (*p*_*Y RI*,0_ = 0.85). This result aligns with previously published work(Cheng *et al*., 2009; Baharian *et al*., 2016), where African-American cohorts ascertained from Southern parts of the United States tended to have higher global African ancestry than those from the North. Additionally, this result highlights that switch counts are sensitive to the assumed initial ancestry proportion, a trend also evident in the parameter dynamics analysis (Figure 1). Given fixed recombination rates and population size, higher switch counts may reflect either higher initial ancestry heterozygosity or more recent admixture, underscoring the identifiably tradeoff inherent in switch-based summaries.

Finally, our results suggest that switch-based models could serve as calibration tools for admixture dating or estimating ancestry proportions, by providing independent constraints alongside tract-length distributions (Chapman and Thompson, 2002; Palamara et al., 2012; Liang and Nielsen, 2014; Janzen *et al*., 2018; Janzen and Miró Pina, 2022) and linkage-disequilibrium–based methods (Moorjani *et al*., 2011; Loh et al., 2013). Under simplifying assumptions such as uniform recombination, tract-length models imply corresponding expectations for ancestry switch counts, since the number of switches on a chromosome is determined by the underlying tract-length distribution (Chapman and Thompson, 2002; Palamara *et al*., 2012; Janzen et al., 2018; Janzen and Miró Pina, 2022). While most admixture models focus on tract-length distributions, ancestry switches offer a complementary summary statistic that also captures the recombination history of a genome. Since switch counts integrate over both map length and heterozygosity decay, they remain robust even when tract boundaries are uncertain. Integrating switch-based and tract-based frameworks could enhance the precision of demographic inference in admixed populations. More broadly, our framework provides a foundation for quantifying the cumulative impact of recombination on ancestry structure, offering a scalable tool for quantifying how demographic, genetic, and evolutionary forces shape mosaic genomes across species.

An important next step is to extend this framework, which currently models the generation of new switches, to include the dynamics of retained switches over time. Tracking the persistence and visibility of junctions across generations would connect switch accumulation directly to the observable ancestry mosaic, providing a bridge between theoretical expectations and local ancestry inference. This extension could also yield probabilistic estimates of switch locations, improving the accuracy of HMM-based local ancestry models. Future versions of the model could additionally incorporate demographic complexity, including migration, time-varying effective population sizes, and multi-way admixture histories, enabling switch-based inference to capture a broader range of evolutionary scenarios.

### Limitations

The current model is limited to a fixed effective population size with two ancestral populations, though the equations could be generalized to non-constant effective population sizes and more ancestries. Additionally, we assume an idealized population with no selection, a single pulse of admixture without subsequent migration, and limited demographic phenomena. In the empirical applications, population-specific recombination maps were inferred from the ancestral source populations. These maps are used to approximate recombination in admixed individuals, an assumption that is most appropriate over the short post-admixture timescales considered here and at broad genomic scales. If recombination landscapes in admixed individuals deviate substantially from those of the parental populations, both the spatial distribution and total number of ancestry switches may differ from model expectations, which could affect inference. In applications to human data, recombination maps represent population-level averages that integrate over sex-specific and individual-level variation in recombination. These rates reflect the net outcome of multiple biological processes shaping recombination and do not allow non-additive effects in admixed individuals to be partitioned explicitly. In addition, LD-based recombination maps may be noisier in regions of low SNP density, and we did not explicitly model uncertainty in recombination rate estimates. Because switch accumulation reflects recombination integrated across extended genomic regions, it is expected to be less sensitive to fine-scale variation in local map estimates. When using the model in another species or to study hybridization, one should carefully consider whether recombination in the parental population is predictive of recombination in the admixed population as this varies across species (Salomé *et al*., 2012; Venu et al., 2024). Lastly, our equation is focused on describing the expectation of the number of newly generated junctions. Thus, we have not modeled the transmission of junctions from parent to offspring.

### Conclusion

Our study provides a unified theoretical framework for understanding how recombination shapes ancestry patterns after admixture. By linking the expected number of ancestry switches to recombination rate, ancestry heterozygosity, and effective population size, we extend classical junction theory to reflect realistic recombination landscapes and population histories. The close agreement among theoretical predictions, forward simulations, and empirical data demonstrates that ancestry switches encode a predictable signature of population history. Together, these results position ancestry switches as a fundamental and interpretable measure of how recombination reshapes genomic ancestry.

## Acknowledgments

The authors would like to thank members of the Edge and Pennell labs for helpful discussions of this work throughout the process. SN, ALK, and JAM were supported by NIH grant R35GM159982. RAP was supported by NIH grant R35GM137758. JAM was also supported by the WiSE Gabilan Assistant Professorship at USC.

## Web Resources

Code and data for analytical equations and simulations: https://github.com/ShirNat/Quantifying-cumulative-ancestry-switches-in-an-admixed-genome

1000 Genomes NYGC data: https://ftp.1000genomes.ebi.ac.uk/vol1/ftp/data_collections/1000G_2504_high_coverage/working/20201028_3202_phased

1000 Genomes strict mask: https://ftp.1000genomes.ebi.ac.uk/vol1/ftp/data_collections/1000_genomes_project/working/20160622_genome_mask_GRCh38/StrictMask/

ENCODE exclusion list regions: https://www.encodeproject.org/annotations/ENCSR636HFF/

FLARE: https://github.com/browning-lab/flare

PLINK 2.0: https://www.cog-genomics.org/plink/2.0/

