## SupplementaryFigsAndText for "Analytical expectations for ancestry junction accumulation in admixed genomes"

### Supplemental Theory

#### Temporal dynamics of expected ancestry switches

To convert our per-generation switch expectation (Equation (1) ) into a fully time–dependent prediction, we must model how local ancestry heterozygosity evolves post-admixture. We do this by tracking the first two moments of the ancestry proportion at a randomly chosen locus, $\mathbb{E}[P_{a_1,g}]$ and $\mathbb{E}[P^2_{a_1,g}]$, over time.

Under neutral Wright-Fisher sampling, the proportion of ancestry at a given locus is a martingale, so its mean remains fixed at the initial admixture level: $\mathbb{E}[P_{a_1,g}] = \mathbb{E}[P_{a_1,0}] = p_{a_1,0}$. However, each generation, when $2N_e$ gametes are sampled to form the next generation, random drift steadily increases the variance of $P_{a_1,g}$. For any random variable $P$, $\mathbb{E}[P^2] = \mathrm{Var}[P] + (\mathbb{E}[P])^2$ . Applying this identity to $P_{a_1,g}$ implies that $\mathbb{E}[P^2_{a_1,g}]$ also increases over time.

Therefore, we can rewrite Equation 1 as

$$\begin{aligned}
\mathbb{E}[S_{L,g}] &= L(r_{a_1} + r_{a_2}) \cdot (\mathbb{E}[P_{a_1,g}] - \mathbb{E}[P^2_{a_1,g}]) \\
&= L(r_{a_1} + r_{a_2}) \cdot (p_{a_1,0} - (\mathrm{Var}[P_{a_1,g}] + (\mathbb{E}[P_{a_1,g}])^2)) \\
&= L(r_{a_1} + r_{a_2}) \cdot (p_{a_1,0} - (\mathrm{Var}[P_{a_1,g}] + p^2_{a_1,0})) \\
&= L(r_{a_1} + r_{a_2}) \cdot (p_{a_1,0}(1 - p_{a_1,0}) - \mathrm{Var}[P_{a_1,g}])
\end{aligned}$$

Having rewritten the switch expectation in terms of $\mathrm{Var}[P_{a_1,g}]$, we now need an explicit recursion for that variance over time. Under Wright–Fisher sampling, fluctuations in $P_{a_1,g}$ come from binomial sampling noise each generation. Concretely, let $X$ be the number of $a_1$ alleles among the $2N_e$ gametes, so that $X \sim \mathrm{Binomial}(2N_e, P_{a_1,g}) \quad \Rightarrow \quad P_{a_1,g+1} = \frac{X}{2N_e}$. Then, by the law of total variance:

$$\mathrm{Var}[P_{a_1,g+1}] = \mathbb{E}[\mathrm{Var}[P_{a_1,g+1}|P_{a_1,g}]] + \mathrm{Var}[\mathbb{E}[P_{a_1,g+1}|P_{a_1,g}]]$$

We evaluate each term in turn:

1. $\mathbb{E}[\mathrm{Var}[P_{a_1,g+1}|P_{a_1,g}]]$

We first calculate the conditional variance:

$$\begin{aligned}
\mathrm{Var}[P_{a_1,g+1}|P_{a_1,g}] &= \mathrm{Var}[\frac{X}{2N_e}] \\
&= \frac{1}{(2N_e)^2} \cdot \mathrm{Var}[X] \\
&= \frac{1}{(2N_e)^2} \cdot 2N_e \cdot P_{a_1,g}(1 - P_{a_1,g}) \\
&= \frac{P_{a_1,g}(1 - P_{a_1,g})}{2N_e}
\end{aligned}$$

Then, we take the expectation and obtain:

$$\begin{aligned}
\mathbb{E}[\frac{P_{a_1,g}(1 - P_{a_1,g})}{2N_e}] &= \frac{1}{2N_e}(\mathbb{E}[P_{a_1,g}] - \mathbb{E}[P^2_{a_1,g}]) \\
&= \frac{1}{2N_e}(\mathbb{E}[P_{a_1,g}] - (\mathrm{Var}[P_{a_1,g}] + (\mathbb{E}[P_{a_1,g}])^2)) \\
&= \frac{1}{2N_e}(\mathbb{E}[P_{a_1,g}] \cdot (1 - \mathbb{E}[P_{a_1,g}]) - \mathrm{Var}[P_{a_1,g}]) \\
&= \frac{p_{a_1,0}(1 - p_{a_1,0}) - \mathrm{Var}[P_{a_1,g}]}{2N_e}
\end{aligned}$$

2. $\mathrm{Var}[\mathbb{E}[P_{a_1,g+1}|P_{a_1,g}]]$

   Given that the binomial is unbiased, the conditional expectation is:

$$\mathbb{E}[P_{a_1,g+1}|P_{a_1,g}] = P_{a_1,g}$$

   Next, we take the variance and obtain:

$$\mathrm{Var}[P_{a_1,g}]$$

Finally, putting the terms together we get:

$$\begin{aligned}
\mathrm{Var}[P_{a_1,g+1}] &= \mathbb{E}[\mathrm{Var}[P_{a_1,g+1}|P_{a_1,g}]] + \mathrm{Var}[\mathbb{E}[P_{a_1,g+1}|P_{a_1,g}]] \\
&= \frac{p_{a_1,0}(1 - p_{a_1,0}) - \mathrm{Var}[P_{a_1,g}]}{2N_e} + \mathrm{Var}[P_{a_1,g}] \\
&= \frac{p_{a_1,0}(1 - p_{a_1,0})}{2N_e} + \mathrm{Var}[P_{a_1,g}](1 - \frac{1}{2N_e})
\end{aligned}$$

For notational convenience, define:

$$V_g = \mathrm{Var}[P_{a_1,g}], \quad a = 1 - \frac{1}{2N_e}, \quad \text{and} \quad b = \frac{p_{a_1,0}(1-p_{a_1,0})}{2N_e}$$

The variance recursion then becomes the standard non-homogeneous first-order linear recurrence,

$$V_{g+1} = aV_g + b$$

with the general solution:

$$V_g = V_0 \cdot a^g + b \cdot \sum_{k=0}^{g-1} a^k$$

By construction, generation 0 corresponds to the instant of admixture, at which we assume the local ancestry proportion is exactly $p_{a_1,0}$ at every locus (no sampling variance), so that $V_0 = 0$. Thus, only the summation remains. This summation contains a geometric series $\sum_{k=0}^{g-1} a^k = \frac{1-a^g}{1-a}$, such that

$$\begin{aligned} V_g &= b \cdot \frac{1-a^g}{1-a} \\ &= \frac{p_{a_1,0}(1-p_{a_1,0})}{2N_e} \cdot \frac{1-\left(1-\frac{1}{2N_e}\right)^g}{1-\left(1-\frac{1}{2N_e}\right)} \\ &= p_{a_1,0}(1-p_{a_1,0}) \cdot \left(1-\left(1-\frac{1}{2N_e}\right)^g\right) \end{aligned}$$

Finally, we can substitute $V_g = \mathrm{Var}[P_{a_1,g}]$, into Equation (1)

$$\begin{aligned} \mathbb{E}[S_{L,g}] &= L(r_{a_1}+r_{a_2}) \cdot (p_{a_1,0}(1-p_{a_1,0}) - \mathrm{Var}[P_{a_1,g}]) \\ &= L(r_{a_1}+r_{a_2}) \cdot \left(p_{a_1,0}(1-p_{a_1,0}) - p_{a_1,0}(1-p_{a_1,0}) \cdot \left(1-\left(1-\frac{1}{2N_e}\right)^g\right)\right) \\ &= L(r_{a_1}+r_{a_2}) \cdot p_{a_1,0}(1-p_{a_1,0}) \cdot \left(1-\left(1-\left(1-\frac{1}{2N_e}\right)^g\right)\right) \\ &= L(r_{a_1}+r_{a_2}) \cdot p_{a_1,0}(1-p_{a_1,0}) \cdot \left(1-\frac{1}{2N_e}\right)^g \end{aligned} \tag{2}$$

This explicit derivation confirms that the expected number of ancestry switches in generation $g$ decays geometrically with factor $(1-\frac{1}{2N_e})^g$, linking junction formation directly to the erosion of ancestry heterozygosity under drift.

This is the same as Equation (2) from the *main text*.

### Supplemental Material and Methods

#### Quantifying ancestry switches via tree sequence outputs

**Algorithm S1:** Switch-counting algorithm

**Input:** Tree sequence at generation $G$ with founders remembered at the time of admixture; genomic segment $[0, L)$

**Output:** $\bar{s}$ = mean number of ancestry switches per haplotype in the admixed population

**Function** `AncestryLabel(`$h, I$`)`:
  `// Traverse upward from the node for haplotype` $h$ `in the marginal tree spanning interval` $I$ `until the first remembered ancestor is reached.`
  `// Read that ancestor's founding-ancestry label (`$a_1$ `or` $a_2$`)`
  **return** $\in \{a_1, a_2\}$

$H \leftarrow$ set of sampled haplotype nodes from the admixed population
**for** $h \in H$ **do**
  $s_h \leftarrow 0$
  **for** *each pair of adjacent intervals* $(I_k, I_{k+1})$ *spanning* $[0, L)$ **do**
    $a_k \leftarrow$ `AncestryLabel`$(h, I_k)$
    $a_{k+1} \leftarrow$ `AncestryLabel`$(h, I_{k+1})$
    **if** $a_k \neq a_{k+1}$ **then**
      $s_h \leftarrow s_h + 1$

$\bar{s} \leftarrow \frac{1}{|H|} \sum_{h \in H} s_h$
`// The switches are averaged per haplotype`
**return** $\bar{s}$

### Supplemental Figures and Tables

#### Parameter Dynamics

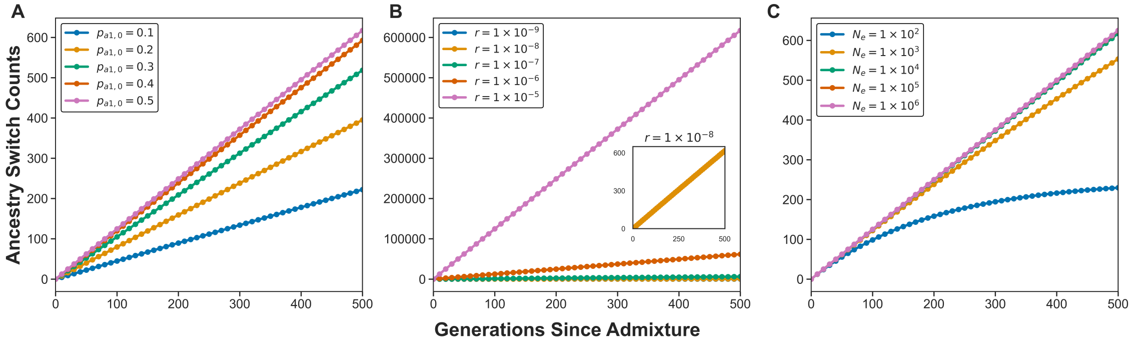


Figure S1: Cumulative theoretical ancestry switch counts after $G$ generations under varying model parameters. For panel A) we vary ancestry proportion $p_{a1,0}$, B) we vary recombination rate $r$, and C) we vary effective population size $N_e$. The total expected new switch counts are calculated from Equation (3) for a haploid chromosome of length $L = 2.5 \times 10^8$ bp. Baseline parameters $r_{a_1} = r_{a_2} = 10^{-8}$, $p_{a_1,0} = 0.5$, and $N_e = 10^4$; values shown for $G \in \{0, 100, \ldots, 500\}$. All expectations are per haplotype.

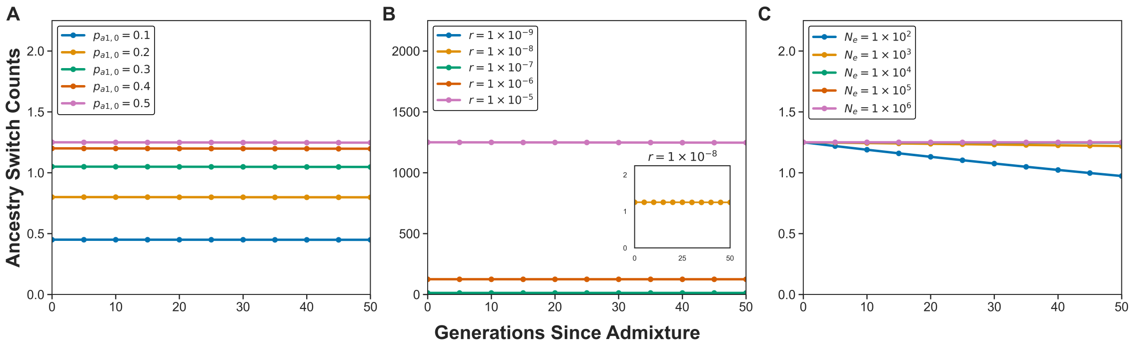


Figure S2: Theoretical per-generation ancestry switch counts under varying model parameters. For panel A) we vary ancestry proportion $p_{a1,0}$, B) we vary recombination rate $r$, and C) we vary effective population size $N_e$. The per-generation switch counts are calculated using Equation (2) for a haploid chromosome of length $L = 2.5 \times 10^8$ bp. Baseline parameters $r_{a_1} = r_{a_2} = 10^{-8}$, $p_{a_1,0} = 0.5$, and $N_e = 10^4$; values shown for $G \in \{0, 10, \ldots, 50\}$. All expectations are per haplotype.

#### Simulation framework and ancestry switch estimation

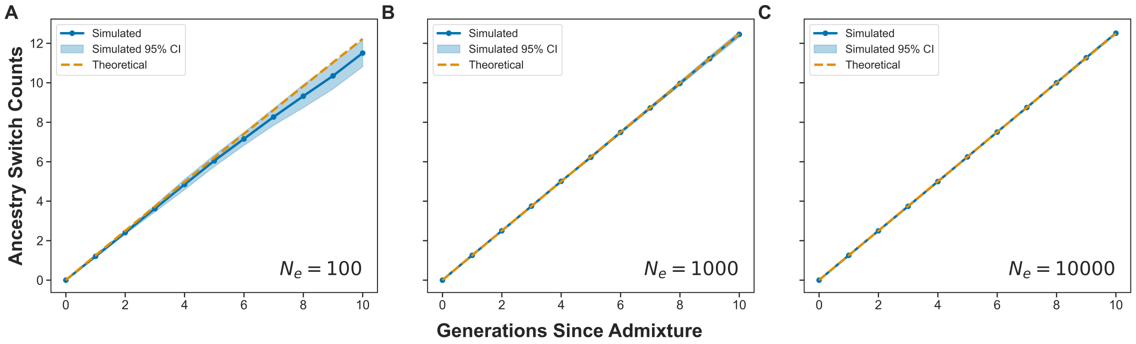


Figure S3: Comparison of simulated and theoretical ancestry switch counts under constant recombination scenario across different population sizes. The mean number of ancestry switches observed across 10 forward-time simulation replicates (blue line) with 95% confidence intervals (shaded region)is compared with theoretical expectations (dashed orange line). Simulated values were obtained from TreeSeq outputs of SLiM simulations, and theoretical expectations were calculated using Equation (3). Results are shown for populations with effective size A) $N_e = 100$, B) $N_e = 1000$, and C) $N_e = 10000$.

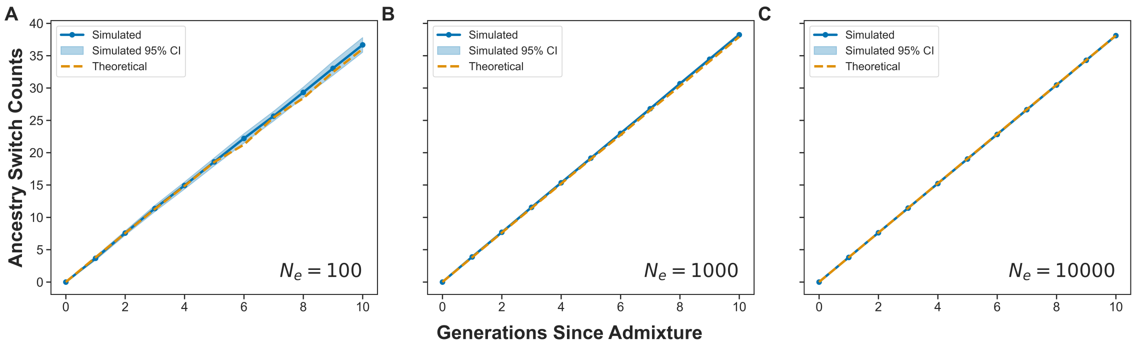


Figure S4: Comparison of simulated and theoretical ancestry switch counts under variable recombination scenario across different population sizes. Recombination rates were drawn from a generated recombination map. The mean number of ancestry switches observed across 10 forward-time simulation replicates (blue line) with 95% confidence intervals (shaded region)is compared with theoretical expectations (dashed orange line). Simulated values were obtained from TreeSeq outputs of SLiM simulations, and theoretical expectations were calculated using Equation (4b). Results are shown for populations with effective size A) $N_e = 100$, B) $N_e = 1000$, and C) $N_e = 10000$.

Table S1: Empirical and theoretical ancestry switch counts per chromosome in African Americans 14 generations post-admixture. Theoretical expectations are computed using full chromosome lengths taken from the hg38 genome assembly, whereas empirical estimates are based on filtered (callable) chromosome lengths after removing problematic genomic regions identified by ENCODE (see *Methods*). Empirical switch counts are generally lower than theoretical expectations. The parameter $p_0$ denotes the initial proportion of African ancestry at admixture. Empirical values are reported as point estimates with 95% confidence intervals (lower–upper).

| Chromosome | Length used in theoretical count (Mb) | Theoretical switches ($p_0 = 0.75$) | Theoretical switches ($p_0 = 0.85$) | Length used in empirical count (Mb) | Empirical switches (95% CI) |
|---|---|---|---|---|---|
| 1 | 248.96 | 9.63 | 6.55 | 179.19 | 6.142 (5.213–7.117) |
| 2 | 242.19 | 9.24 | 6.28 | 195.99 | 4.880 (4.159–5.790) |
| 3 | 198.30 | 7.70 | 5.24 | 163.27 | 4.236 (3.548–5.007) |
| 4 | 190.21 | 8.32 | 5.65 | 156.41 | 3.539 (2.881–4.292) |
| 5 | 181.54 | 7.11 | 4.83 | 146.36 | 4.013 (3.252–4.830) |
| 6 | 170.81 | 6.79 | 4.62 | 139.27 | 3.394 (2.837–3.990) |
| 7 | 159.35 | 6.79 | 4.62 | 121.65 | 3.214 (2.720–3.780) |
| 8 | 145.14 | 7.06 | 4.80 | 118.89 | 2.659 (2.156–3.250) |
| 9 | 138.39 | 7.65 | 5.20 | 89.95 | 3.254 (2.745–3.739) |
| 10 | 133.80 | 6.20 | 4.21 | 106.11 | 3.114 (2.620–3.641) |
| 11 | 135.09 | 5.64 | 3.83 | 107.13 | 2.946 (2.530–3.517) |
| 12 | 133.28 | 5.66 | 3.85 | 106.13 | 3.113 (2.635–3.680) |
| 13 | 114.36 | 4.35 | 2.96 | 81.48 | 2.342 (1.837–2.901) |
| 14 | 107.04 | 4.19 | 2.85 | 71.76 | 1.604 (1.239–2.032) |
| 15 | 101.99 | 4.76 | 3.24 | 62.38 | 1.651 (1.266–2.030) |
| 16 | 90.34 | 4.99 | 3.39 | 57.60 | 1.129 (0.842–1.448) |
| 17 | 83.26 | 6.99 | 4.75 | 56.85 | 1.458 (1.135–1.823) |
| 18 | 80.37 | 8.28 | 5.63 | 63.26 | 1.082 (0.837–1.378) |
| 19 | 58.62 | 4.16 | 2.83 | 37.19 | 0.697 (0.510–0.885) |
| 20 | 64.44 | 3.65 | 2.48 | 49.27 | 0.843 (0.646–1.083) |
| 21 | 46.71 | 3.03 | 2.06 | 27.21 | 0.632 (0.415–0.830) |
| 22 | 50.82 | 3.17 | 2.15 | 25.23 | 0.702 (0.500–0.917) |
